# Identification of protein secretion systems in bacterial genomes using MacSyFinder version 2

**DOI:** 10.1101/2023.01.06.522999

**Authors:** Sophie S Abby, Rémi Denise, Eduardo PC Rocha

## Abstract

Protein secretion systems are complex molecular machineries that translocate proteins through the outer membrane and sometimes through multiple other barriers. They have evolved by co-option of components from other envelope-associated cellular machineries, making them sometimes difficult to identify and discriminate. Here, we describe how to identify protein secretion systems in bacterial genomes using the MacSyFinder program. This flexible computational tool uses the knowledge gathered from experimental studies to identify homologous systems in genome data. It can be used with a set of pre-defined MacSyFinder models—”TXSScan”, to identify all major secretion systems of diderm bacteria (*i*.*e*., with inner and LPS-containing outer membranes) as well as evolutionarily related cell appendages (pili and flagella). For this, it identifies and clusters co-localized genes encoding proteins of secretion systems using sequence similarity search with Hidden Markov Model (HMM) protein profiles. Finally, it checks if the clusters’ genetic content and genomic organization satisfy the constraints of the model. TXSScan models can be altered in the command line or customized to search for variants of known secretion systems. Models can also be built from scratch to identify novel systems. In this chapter, we describe a complete pipeline of analysis, starting from i) the integration of information from a reference set of experimentally studied systems, ii) the identification of conserved proteins and the construction of their HMM protein profiles, iii) the definition and optimization of “macsy-models”, and iv) their use and online distribution as tools to search genomic data for secretion systems of interest. MacSyFinder is available here: https://github.com/gem-pasteur/macsyfinder, and MacSyFinder models here: https://github.com/macsy-models.

## 1. Introduction

Bacteria produce proteins to interact with other individuals, prokaryotes or eukaryotes, to operate changes in their local environment, or to uptake resources. Many of the proteins involved in these processes need to be secreted to the outside of the cell. Bacteria with an LPS-containing outer membrane (henceforth called diderms) face a formidable challenge to secrete these proteins because they must transport them through the inner membrane, the cell wall, the outer membrane, and sometimes additional extra barriers such as the bacterial capsule and membranes of other cells. The complexity of these molecular processes and the key roles of protein secretion systems in bacterial ecology and virulence have spurred much interest in their study (for recent reviews, see [1–9]). There are at least eight well-known protein secretion systems in diderms, and now even up to ten described (numbered from T1SS to T11SS, excluding the T7SS from Mycobacteria), but others probably remain to be uncovered [10].

There are few computational tools to identify and characterize protein secretion systems in bacterial genomes (for a list see Table 1 of [10]). Their development becomes urgent in light of the availability of many thousands of genomes and the ease with which new ones can be sequenced. These tools should be able to identify components of the protein secretion systems and assess if they are sufficient to define an instance of a given system. When the components are highly conserved proteins, they can be identified with high sensitivity by sequence similarity search. The identification of fast-evolving components might be more complicated because of poor sequence conservation. Additionally, some components may not be strictly necessary for a functional system, and it may be difficult to know which of the two factors explains their absence from an instance of the system. Under these conditions, it is useful to split the components of secretion systems into those that should be present in the instance (“mandatory”) and those that may be absent (“accessory”). The former correspond to highly conserved, easily identifiable components, while the latter correspond to components that may be lacking in systems because they are missing or not detected. This nomenclature does not presume anything about the biological role of the accessory components: they may be biologically essential but unidentifiable by sequence similarity search. This classification aims at describing the system in a way that facilitates its identification in genomes. We will use it throughout this text.

**Table 1.** Useful online resources to design macsy-models for the detection of secretion systems.

| Resource [reference] | Type | Commands |
| --- | --- | --- |
| NCBI/Blast [40] | Sequence database indexing<br>Sequence similarity search | makeblastdb<br>blastp |
| Silix [30] | Sequence clustering | silix |
| MAFFT [25] | Multiple sequence alignment | mafft |
| Muscle [26] | Multiple sequence alignment | muscle |
| Seaview [27] | Sequence alignment, edition, and phylogeny | seaview |
| Jalview [28] | Sequence alignment, edition, and analysis | - |
| HMMER [24] | Build HMM profiles and use them for sequence similarity searches | hmmbuild<br>hmmsearch |
| PFAM [41] | Database of HMM protein profiles | - |
| TIGRFAM [42] | Database of HMM protein profiles | - |
| InterProScan [43] | Identify conserved domains using several resources | - |
| MacSyFinder [18, 19] | Macromolecular systems detection from models of systems | macsyfinder |
| TXSScan [10, 12, 19] | MacSyFinder-based package of models (or "macsy-model") to detect T1SS-T9SS and related appendages (Type IV filament super-family pili and bacterial flagellum) | macsyfinder |
| TXSScan@Galaxy/Pasteur | Enables to run online and interactively MacSyFinder with TXSScan on user uploaded genomes | - |
| TXSSdb [10] | Database of secretion systems detected with TXSScan in 1528 genomes of diderm bacteria | - |

The evolution of secretion systems has involved the co-option of many components from other molecular machineries [11]. These components have sometimes been co-opted in turn for other cellular machineries [12]. As a result, many components of protein secretion systems have homologs in other systems [13, 14]. This increases the risk of misidentifications. For example, the T3SS and the flagellum are evolutionarily related, and several of their core components belong to homologous families [13]. In this specific case, the discrimination between the two systems is facilitated by the existence of flagellum-associated mandatory proteins that are always absent from T3SS (*e*.*g*., FlgB) and vice-versa (*e*.*g*., the secretin). These components can be qualified as “forbidden” in the other system to prevent misidentifications.

The analysis of the genetic context can also improve the discrimination between protein secretion systems and other molecular systems. For example, the components of the T3SS are usually encoded in one single locus, which facilitates their discrimination from those of the flagellum [15]. Another example is provided by the three mandatory components of the T1SS, which all have homologs in other systems [16], even if none of the other systems includes all three [17]. Notably, the abc (ABC-transporter) and the mfp (membrane fusion protein) components are systematically encoded in the same locus in T1SS (and only in this system). Hence, three types of information facilitate the unambiguous detection of secretion systems: identifying pertinent and forbidden components, the completeness of the set of components, and their genetic organization. This information can be put together in a model of the system that can be used by a computer program that we developed and called MacSyFinder for “Macromolecular System Finder” [18]. This program, now in its second version (“v2”), searches genomes for instances satisfying the characteristics described in the model in terms of genetic content (or “quorum”) and organization [18, 19].

Sometimes good computational models of protein secretion systems are not available. Creating novel (or better) models requires to identify the relevant components and their genetic organization. Most secretion systems were studied on a small number of bacteria, where they were sometimes remarkably well characterized in terms of their components, their genetic regulation, their structure, and sometimes their assembly pathways. In contrast, the other instances of the systems are usually very poorly characterized. The challenge posed to the researcher interested in identifying novel instances of a given type of system is thus to produce models with relevant descriptions of the current knowledge of the system. This is difficult because the number of components and their organization may vary widely. For example, the Type IV secretion system (T4SS) locus of *Legionella pneumoniae* is encoded by more than twice the number of genes of the *vir* T4SS of *Agrobacterium tumefaciens* [20, 21]. In addition, most T4SS are encoded in a single locus, but there are intracellular pathogens where they are encoded in several distant loci [22]. The key point in the production of novel models is thus the identification of the traits that are conserved and can be most useful to identify a certain type of systems.

The production of models involves generalizing knowledge that was obtained from specific examples. These models are quantitative representations of the composition and organization of the known systems. When they work, they vastly facilitate the identification of homologous systems. When they fail, they highlight gaps in our understanding of the system, which often raises interesting biological questions.

This text shows how one can use MacSyFinder version 2 (v2) to identify protein secretion systems with the pre-defined package of MacSyFinder models TXSScan (*see* **Figure 1**, section 3.3). These models define the components of the system, the minimal number of mandatory and accessory components (quorum), and their genetic organization. They have been validated and shown to perform well: they identify the vast majority of known systems [10, 12]. An earlier version of TXSScan has been used to identify over 10,000 systems in 1258 genomes of bacteria (available in MacSyDB/TXSSdb, *see* **Table 1**). We recently developed a version adapted to MacSyFinder v2, whose content is displayed in **Figure 1** and **Figure 2**. It includes the best-known protein secretion systems from diderm bacteria as well as the flagellum related to the Type III secretion system (T3SS) and the entire family of pili related to the Type II secretion system (T2SS), namely the Type IV filament super-family (TFF-SF). Yet, these models may be inadequate in certain specific situations. In this chapter, we show how they can be modified or built from scratch to identify novel, or variants of a secretion system. MacSyFinder uses Hidden Markov model (HMM) protein profiles specified in a model to search for components of a system in a file of protein sequences. In a nutshell, it collects clusters of co-localized components in the genome and checks if combinations of these clusters may result in valid systems relative to the gene content and co-localization specified in the systems’ models [19]. MacSyFinder then outputs the results of the protein profile searches and the information about the identified secretion systems.

**Figure 1.**
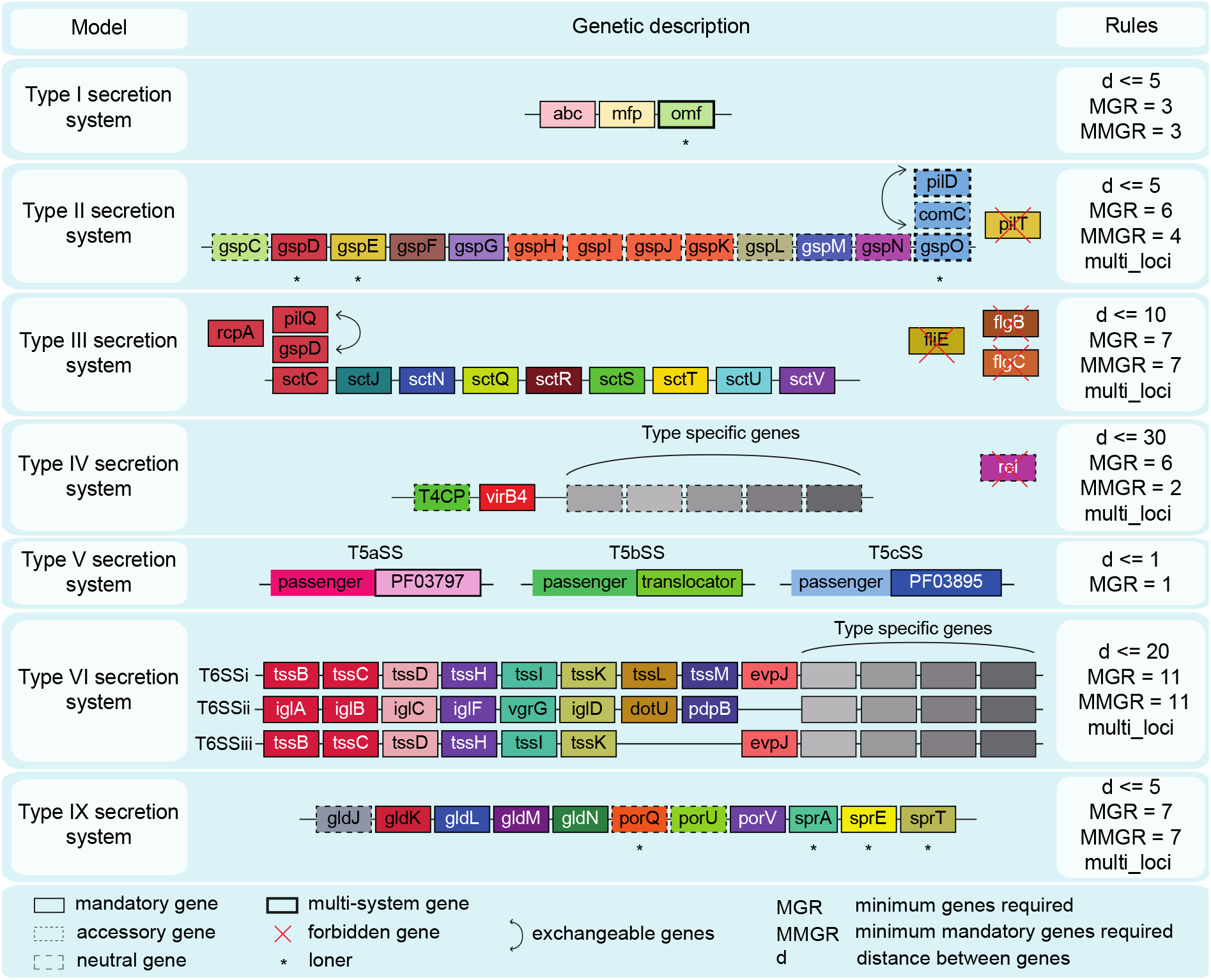
Models of protein secretion systems from diderm bacteria available in TXSScan version 1.1.1. All of these systems are available from TXSScan, in the bacteria/diderm sub-folder. Each box represents a component and its status in the system’s model: “mandatory” (plain), “accessory” (dashed), “neutral” (bracketted) or “forbidden” (red cross). Within a system panel: the families of homologous proteins are represented in columns and colored identically. The minimal number of genes required to reach the quorum and infer the system’s (“MGR” and “MMGR”, see Figure’s legend) is indicated, as well as the co-localization parameter of the system (d). Curved double-headed arrows indicate exchangeable components. Additional features specific to a component can be found in **Table 2**. Figures and legends are freely reproduced with modification from [10, 12] (as specified by the Creative Commons Attribution (CC BY) license version 4.0).

**Figure 2.**
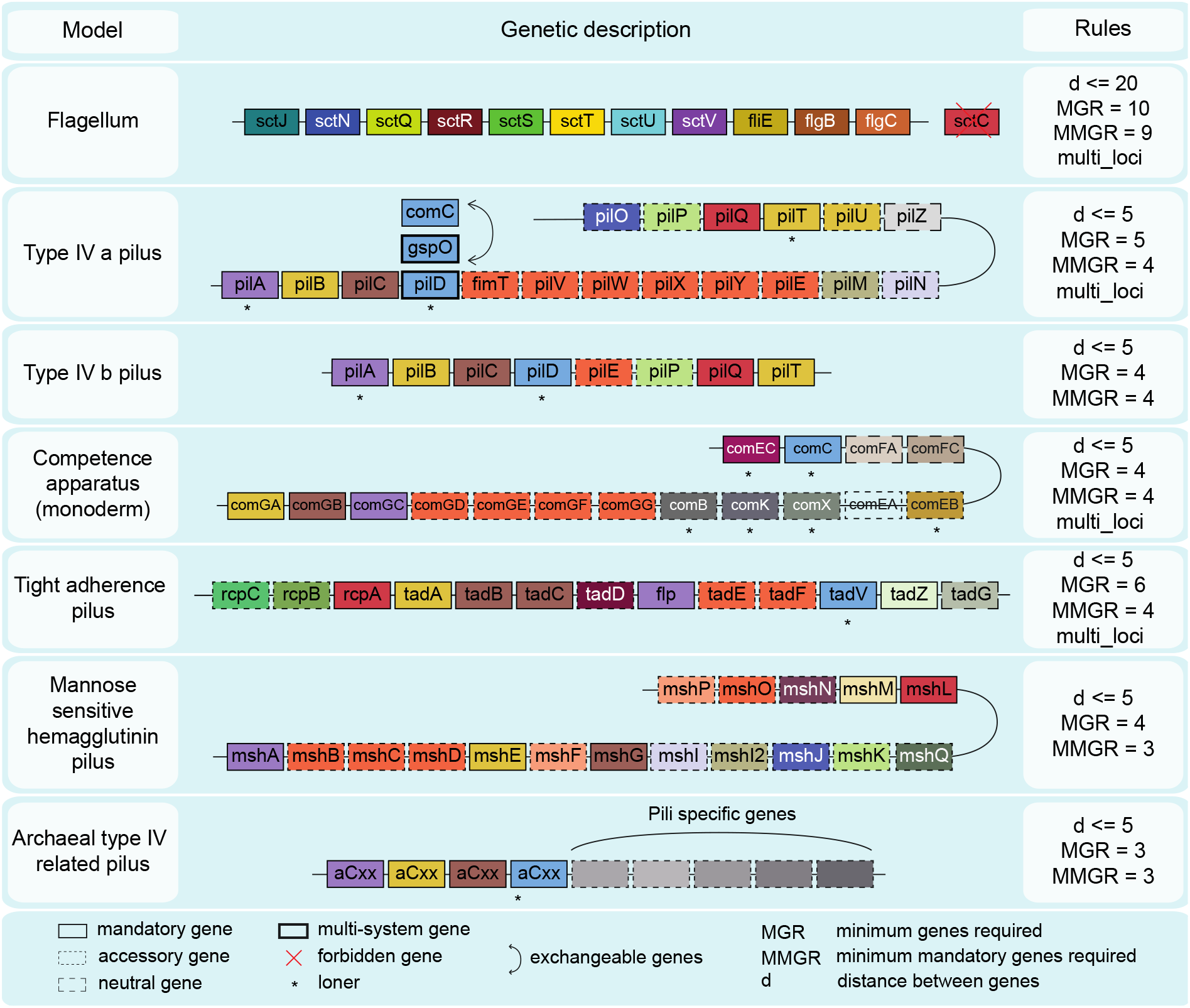
Models of appendages related to protein secretion systems available in TXSScan version 1.1.1. Diverse pili and flagella (the bacterial flagellum at the top of the figure, and the archaeal flagellum, at the bottom of the figure) are related to diderm bacteria secretion systems. Some variants of these systems have also been shown to be involved in protein secretion. The legend is the same as for **Figure 1**.

In the next section, we indicate the data and software required to use and design models for MacSyFinder. Then, in the following section, we describe how to define models and protein profiles to identify protein secretion systems of interest. Finally, we explain how the designed models can be easily shared with the community for reuse using the new *macsydata* tool via the MacSy Models central repository here available: https://github.com/macsy-models.

**Table 2.** Keywords to specify the XML models. For more details, see the MacSyFinder Modeller guide: https://macsyfinder.readthedocs.io/en/latest/modeler_guide/index.html#

| Keyword in XML model definition<br>(Command-line over-riding) | Required? (default) | Level(s) | Description |
| --- | --- | --- | --- |
| model | Yes | upper (first) | keyword needed to start the definition of the system's model |
| inter_gene_max_space<br>(--inter-gene-max-space) | Yes | model, gene | “co-localization” parameter. Defines the maximal number of genes between two consecutive components of the system for a cluster of components to be created |
| min_genes_required<br>(--min-genes-required) | No (number of mandatory genes or min_mandatory_genes_required if declared) | model | minimal number of mandatory and accessory components |
| min_mandatory_genes_required<br>(--min-mandatory-genes-required) | No (number of mandatory genes, or min_mandatory_genes_required if declared) | model | minimal number of mandatory components |
| multi_loci<br>(--multi-loci) | No (false, *i.e.*, "0") | model | whether the system can be encoded on several main loci |
| gene | Yes | second | component of the system (below "model") |
| multi_system | No (false, *i.e.*, "0") | gene | whether the gene can participate in several instances of a same system |
| multi_model | No (false, *i.e.*, "0") | gene | whether the gene can participate in several instances of different models |
| name | Yes | gene | the name of the gene to be used in report files, and base name of the corresponding HMM protein profile |
| loner | No (false, *i.e.*, "0") | gene | whether the gene can be found encoded outside of the system's main loci - *i.e.*, an exception to the co-localization rule defined at the system level |
| exchangeables | No | gene | whether the gene can be replaced by another gene in the list of the components (quorum). The gene can then be exchanged with one of the genes listed at the inferior level. |

## 2. Materials

### 2.1 Sequence data

MacSyFinder analyzes protein sequences to identify protein secretion systems. Therefore, the protein sequences should all be stored in one single file in FASTA format. This file represents one of several types of information, which must be specified by the “--db-type” option:

- When the proteins are from one single genome (or even possibly diverse sources) and the corresponding genes’ relative order is unknown, the file type is “unordered”. In this case, the program can identify components and check if the components’ quorum is respected. It cannot, however, check the genetic organization of the model.
- When proteins are from one single genome (or replicon, *i*.*e*., chromosome or plasmid) and are ordered following the position of genes in the genome, then the file type is “ordered_replicon”. The “ordered_replicon” mode allows the use of all the available criteria (quorum of components and genetic organization) to identify instances of the system, and as such, is the most powerful. This mode should be used to create, test, and validate a new model and the corresponding protein profiles. It will be the focus of this protocol.
- The type “gembase” is similar to the “ordered_replicon” but has special identifiers that allow the analysis of multiple “ordered” genomes in one single step (see MacSyFinder’s documentation).

### 2.2 Pre-defined models available in TXSScan

TXSScan is a set of pre-defined MacSyFinder models and profiles (also called “macsy-model” package), to detect the best-studied protein secretion systems in diderms (*see* **Figure 1**, with the T1SS, T2SS, T3SS, T4SS, T5SS, T6SS, T9SS) and related appendages with homologs in diderm and monoderm bacteria, and even in archaea (*see* **Figure 2**, with the bacterial flagellum and the pili members of the type IV filament super-family) [10, 12, 19]. It has to be noted that since MacSyFinder version 2, it is possible to class MacSyFinder models in sub-folders with a hierarchy enabling to run MacSyFinder only on a subset of relevant models at once. For TXSScan, the hierarchy of models follows the split in kingdoms (bacteria versus archaea), then in membrane types (monoderm versus diderm within bacteria). TXSScan models are used as examples throughout the following sections. The files of TXSScan (HMM profiles and model files) can easily be installed for MacSyFinder using the *macsydata* tool from the MacSyFinder suite (*see* section 3.3.1). Were they to be created/modified by the user, they should be copied from the GitHub repository (https://github.com/macsy-models/TXSScan/tags) and placed at a recognizable location to be used with MacSyFinder (*see* section 3.3.1). MacSyFinder can also be used online with the TXSScan models (*see* **Table 1**). Currently, only the standalone version of MacSyFinder allows the modification of the TXSScan models and the introduction of novel protein profiles.

### 2.3 Software

We list the resources of interest for this protocol in **Table 1**. To run MacSyFinder, one needs to install HMMER (version >3.0) and MacSyFinder [19, 23, 24]. The latter requires a Python interpreter (version >3.6) that must be installed beforehand. See MacSyFinder’s online documentation for more details [19].

To build novel HMM protein profiles, one also needs a program to make multiple sequence alignments (*e*.*g*., MAFFT or Muscle [25, 26]), an alignment editor (*e*.*g*., Seaview or Jalview [27, 28]), a sequence similarity search program (*e*.*g*., Blast or DIAMOND [23, 29]) and a program to cluster proteins by sequence similarity (*e*.*g*., Silix or MCL [30, 31]).

## 3. Methods

The procedure to design a novel system’s model follows several steps (*see* **Figure 3**). It starts with identifying the relevant components and their genetic organization from an identified reference dataset of experimentally validated instances of the system. Next, the HMM protein profiles for each component can be built from the reference dataset or retrieved from public databases. These three types of information (components’ list, genetic architecture, and profiles) can then be used to formulate the model. Finally, the model is used to analyze an independent dataset of experimentally validated systems to check they are indeed annotated. At the end of this process, one should be able to use the model to identify novel instances of the system and know the sensitivity of the procedure.

**Figure 3.**
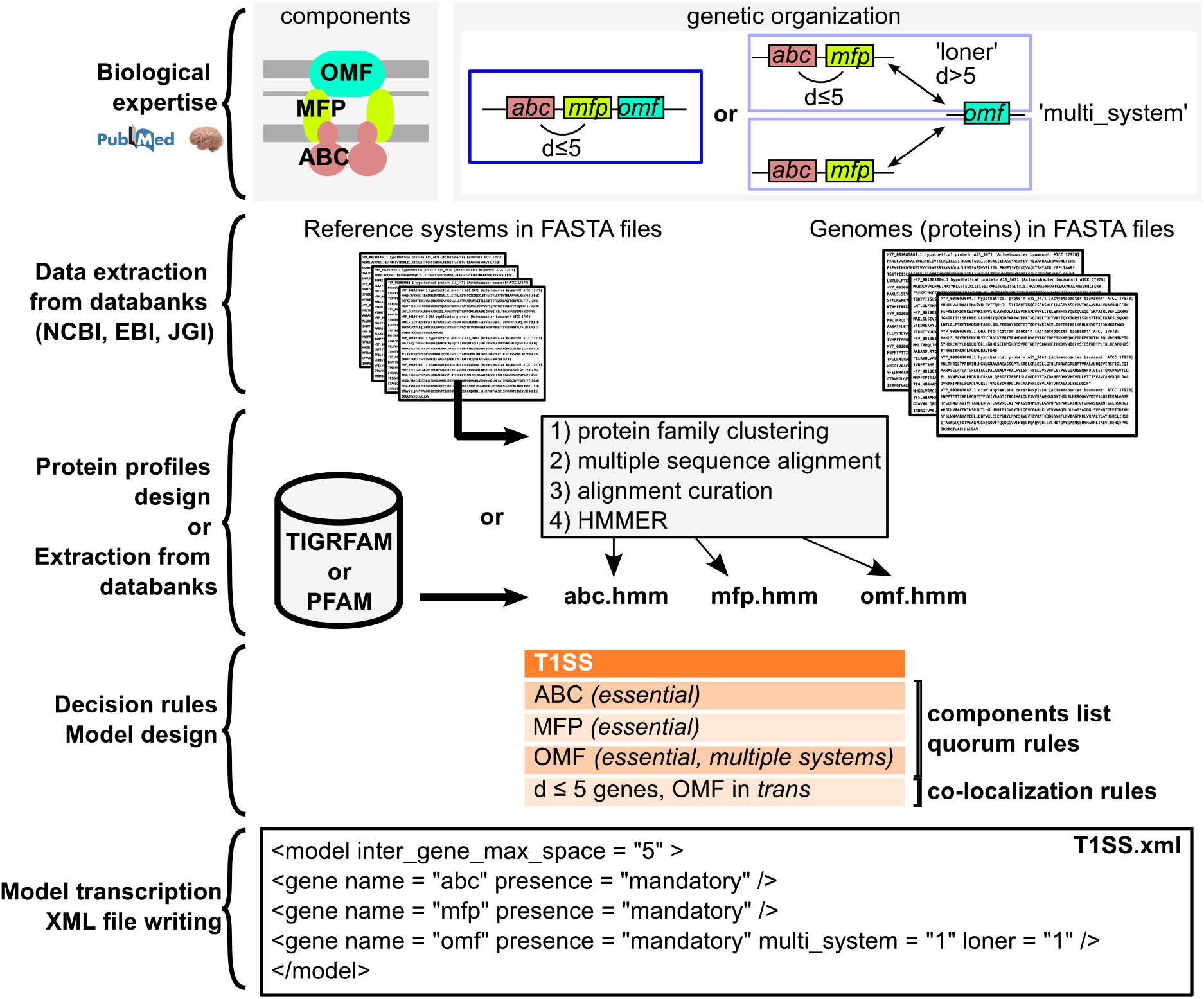
Overview of the protocol to design a macsy-model package.

### 3.1 Compilation of available information on the system

Gathering the current biological knowledge of the system, especially of its experimentally studied instances, may not be straightforward. The genes and proteins of experimentally studied systems are often named differently; sequence similarity searches may thus be necessary to establish which components in a system are homologous to those of other systems. Occasionally, gene fusions and fissions further complicate the identification of families of homologs. Once the relationships of homology between known instances of the system are well established, its key components can be inventoried and qualified (as *mandatory, accessory*, and *forbidden*). One can then characterize the system in terms of its genetic organization and quorum (*i*.*e*., how many components are required for a system, *see* section 3.1.4). The set of experimentally validated protein secretion systems should be split into two independent datasets (*see* **Note 1**). The *reference* dataset is used to characterize the protein secretion systems, build the models, and construct the protein profiles (if required, *see* **Note 2** and **Note 3**). The *validation* dataset is used to validate the final model (*see* section 3.5). There are alternative procedures for systems with few experimentally validated systems (*see* **Note 4**).

#### 3.1.1 Identification and classification of the components of the secretion system

The most frequently encountered components of a secretion system are typically classified as “mandatory”. They may be inferred from the frequency of each protein family in the *reference* dataset or retrieved from the literature (*see* sections 3.1.3 and 3.1.4) (*see* **Note 2**). The other components are classified as “accessory”. MacSyFinder version 2 introduces another class of “neutral” components. This new class may help annotate genes of particular interest that do not count in the quorum (*e*.*g*., new candidate genes). When it is necessary to distinguish a system from other systems with many homologous components, it may be relevant to introduce and classify some genes specific to the other systems as “forbidden”, as for the T2SS (*e*.*g*., PilT) or the T3SS (FlgB, *see* **Figure 1**). **Table 2** lists the features and terms available to describe the systems’ model.

#### 3.1.2 Extraction of HMM protein profiles from databanks

Each component (named “gene” in the model) must be associated with at least one HMM protein profile. Several public databanks of protein profiles can be queried by keyword (*e*.*g*., the name of the gene) or by sequence for a match to the component (*see* **Table 1**). The InterPro portal is a good starting point for this task since it allows to obtain information on protein profiles from many databanks, including PANTHER, PFAM, SUPERFAMILY, and TIGRFAM [32]. Interestingly, many of these profiles include a threshold score for hit inclusion, such as the “GA” (gathering) scores (*see* **Note 5**). These GA thresholds for bit-scores allow to impose profile-specific criteria to filter the hits. We explain in **Note 5** how the user can compute such a score for its own HMM profiles to enable component-specific filtering (available since MacSyFinder v2). These GA scores can be used to overrule the overall filtering criteria set with the “--i-evalue-sel” and “--coverage-profile” options of *macsyfinder*.

At the end of this procedure, one often has many profiles for some components and none for others. Therefore, one can pick the best matching profile for each component (*see* **Note 1**). However, sometimes there is no profile better than the others. This occurs when profiles match only certain sub-families of the components, *e*.*g*., because they are divergent in sequence. In these cases, it is possible to specify multiple profiles to identify the same component (*see* **Note 6**).

The user will have to build a novel protein profile in one of three typical situations: (1) when none is available in the databases; (2) when one profile can replace a large set of very specific profiles; (3) when one wishes to build profiles that are specific to a system because existing profiles also match components from other types of systems. The new profile should be built using the sequences of the *reference* dataset (*see* also **Note 1** and **Note 3**).

#### 3.1.3 Establishing the model of the genetic architecture

By default, MacSyFinder searches for clusters of co-localized genes encoding components of the system (for ordered datasets of type “ordered_replicon” or “gembase, *see* section 2.1). The maximal distance allowed between consecutive components (“inter_gene_max_space”) can be the same for all the components, or specific to a component. The distance is measured in genes unit, *i*.*e*., a distance of three means that there can be up to three genes between two consecutive components of the system. This distance can be inferred from the *reference* dataset or the secretion systems identified in bacterial genomes (*see* **Note 7**). Additionally, one may define some genes as “loner”, in which case the model allows them to be encoded outside the clusters of co-localized genes (section 3.2.1).

The overall genetic architecture of a system can be specified through the “multi_loci” attribute. The default value (False or “0”) indicates that the system is encoded in a single locus (except for the “loner” components). In contrast, the alternative (True or “1”) authorizes the existence of multiple clusters for an instance of the system. For example, this option is helpful to describe the type IVa pilus or the T9SS since they are often encoded in several loci [10].

#### 3.1.4 Defining the quorum of components

The quorum of the model is the minimal number of components required to validate an instance of the system. It is defined by two parameters: the minimal number of mandatory components (“min_mandatory_genes_required”) and the minimal number of mandatory and accessory components (“min_genes_required”). The values of both parameters are set by default to the number of mandatory components; in this case, the accessory components do not count in the quorum. To make them count, one must specify higher values for the second than for the first parameter. It is particularly important to use prior knowledge to assess the relevance of a locus lacking a mandatory gene.

The quorum can be optimized using the information from the secretion systems identified in bacterial genomes (*see* **Note 8**). Changes in the values of the quorum affect the sensitivity and specificity of the method. Low values authorize the validation of systems with fewer components and less similar to the reference dataset, thus increasing the number of detected systems. This might come at the cost of misidentifications or the validation of non-functional systems. Running the models with higher values of the quorum results in identifying instances that are more similar to the reference dataset, and thus more likely to be true. On the other hand, this results in the identification of fewer instances and may exclude those that are too distinct from the ones in the *reference* dataset.

### 3.2 Formulation of the model

After gathering the available information on the protein secretion system, one must write down the model in the relatively simple XML format pre-defined for MacSyFinder. A list of the keywords of this hierarchical XML grammar is available in **Table 2** and MacSyFinder’s online documentation (*see* the Modeller guide). The grammar used by MacSyFinder was slightly changed and simplified in version 2. Hence, the XML files used with version 1 are not compatible with MacSyFinder v2 (and vice versa).

#### 3.2.1 Defining the model in an XML text file--Example 1: the T1SS

We illustrate the formulation of a simple model with the example of the type I secretion system. This system has three “mandatory” components (**Figure 3**): an ABC transporter (“abc”), a membrane fusion protein (“mfp”), and an outer membrane porin (“omf”). The latter can be co-localized with the two other components (less than 6 genes apart, “inter_gene_max_space” set to 5) or encoded apart in the genome (“loner” attribute set to True or “1”). In addition, omf can be involved in several occurrences of the system (“multi-system” set to True or “1”). The three mandatory components are required to form a full system. The quorum is not specified; it is left to its default value (*i*.*e*., three, the number of mandatory components). The file with the model should be named after the system (“T1SS.xml”). Its content is displayed in **Figure 3**.

#### 3.2.2 Defining the model in an XML text file--Example 2: the T9SS

Larger systems tend to require more complex models. This is well exemplified by the model to identify T9SS (file “T9SS.xml”, https://github.com/macsy-models/TXSScan/blob/bcde1991c027c8d328128599792c80527abdf043/definitions/bacteria/diderm/T9SS.xml).

This model, in line with existing literature [33–36], states that T9SS consist of several mandatory and accessory components encoded in multiple loci (multi_loci = “1”). The quorum allows one “mandatory” and several “accessory” components to be missing. Some components form gene clusters, whereas others are defined as “loners” (**Figure 1**). The SprA component was matched by several profiles of the PFAM and TIGRFAM databanks; we used the “exchangeables” attribute to include them all (*see* also **Note 6**):

~~~
<gene name=“T9SS_sprA_PF14349” presence=“mandatory” loner=“1”><exchangeables><gene name=“T9SS_sprA-2_PF12118” loner=“1”/><gene name=“T9SS_sprA-3_TIGR04189” loner=“1”/> </exchangeables></gene>
~~~

### 3.3 Running MacSyFinder

#### 3.3.1 Organizing the models data or installing the macsy-models from a repository

Since version 2 of MacSyFinder, the macsy-model packages containing required XML models and HMM profiles adopt a pre-defined file architecture. A macsy-model folder must contain one folder gathering the XML model files called “definitions” and one folder called “profiles” containing all the corresponding HMM protein profiles listed as genes’ components in the XML models. Such a file architecture can be automatically created using MacSyFinder’s tool *macsydata* with the “*macsydata init*” sub-command. HMM protein profiles should be in individual files named after the corresponding component. All profiles must have the same file name extension (“.hmm” by default, can be changed with the option “--profile-suffix”). For the T1SS model mentioned above (*see* **Figure 3**), the program requires three HMM files in a directory “profiles”: “abc.hmm”, “mfp.hmm”, and “omf.hmm”.

The path to the macsy-folder directory containing the “definitions” and “profiles” folders must be given in the *macsyfinder* command line through the “--models-dir” option. If the user wants to use predetermined MacSyFinder models, MacSyFinder is shipped since v2 with the companion tool *macsydata* that enables to easily retrieve and install public macsy-models from the dedicated Github “Macsy Models” repository (https://github.com/macsy-models). To install the latest version of the TXSScan macsy-model, one can thus simply run:

~~~
macsydata install TXSScan
~~~

Then all the models available in TXSScan (*see* **Figures 1 and 2**) will be available for search in genomes with the *macsyfinder* command *(see* section 3.3.2).

#### 3.3.2 Identification of the secretion systems

Here we exemplify the procedure to identify instances of T1SS in a protein file of the type “ordered_replicon”, for example, on the proteins from the *Acinetobacter baumannii* ATCC 17978 complete genome downloaded from the NCBI and stored in the “CP000521_proteins.fasta” file [37]. Some example genomes, including the latter, along with different command lines and expected output files, can be found here: https://doi.org/10.6084/m9.figshare.21716426.v1.

The command line to launch the program is:

~~~
macsyfinder --db-type ordered_replicon --replicon-topology circular --sequence-db
CP000521_proteins.fasta -–models TXSScan bacteria/diderm/T1SS
~~~

To search for both the T1SS and T9SS, the command line is the following:

~~~
macsyfinder --db-type ordered_replicon --replicon-topology circular --sequence-db
CP000521_proteins.fasta -–models TXSScan/bacteria/diderm T1SS T9SS
~~~

Or:

~~~
macsyfinder --db-type ordered_replicon --replicon-topology circular --sequence-db
CP000521_proteins.fasta -–models TXSScan bacteria/diderm/T1SS bacteria/diderm/T9SS
~~~

Depending on how the models were developed, the identification of multiple secretion systems can be better done independently, or one can identify them simultaneously. Since the version 2 of MacSyFinder, TXSScan is very efficient at identifying them all at once. For example, to identify all bacterial secretion systems and related appendages available for annotation in TXSScan, one should type:

~~~
macsyfinder --db-type ordered_replicon --replicon-topology circular --sequence-db
CP000521_proteins.fasta -–models TXSScan/bacteria all
~~~

### 3.4 Finding the most relevant information from MacSyFinder’s output files

The results of the detection are printed to files in tabulated text format (“.tsv” for “tabulated separated values”) that can be read by any spreadsheet processing application. They are stored in the MacSyFinder’s output directory (specified using the “-o” option or named automatically as in the examples above). They include configuration and log files, along with the results of MacSyFinder and HMMER. The description of the output files is detailed in MacSyFinder’s documentation.

Some output files are particularly useful for improving the design of a new model. The file “best_solution.tsv” indicates the composition of the identified systems as part of the best solution (maximal set of non-overlapping systems). In contrast, the file “all_systems.tsv” reports all the valid instances of different candidate systems, whether they were selected as part of the best solution or not. The file “rejected_candidates.tsv” reports candidate systems that did not pass the quorum criteria. Together with the files storing raw or filtered HMMER hits (in the “hmmer_results” directory), it gives an overview of the detection process, from the detection of components to the validation of the clusters. For automated downstream analyses of MacSyFinder’s results, the files “best_solution.tsv” and “best_solution_summary.tsv” are particularly interesting. The former contains the components identified in each instance of identified systems with information on the quality of the HMMER matches. The latter describes the content in each type of system for the replicon(s) annotated (*see* **Note 7** and **Note 8**). NB: since version 2 of MacSyFinder the files “results.macsyfinder.json” are not generated anymore since the MacSyView web tool is no longer maintained and distributed.

### 3.5 Optimization, validation, and public sharing of the macsy-models

#### 3.5.1 Optimization

The initial models should be very general to assess the diversity of the instances of the system. They can then be optimized iteratively in function of the results obtained in the analysis of the *reference* dataset (and external data). The output files discussed in section 3.4.2 can be used to identify the parts of the model that must be improved. The specific case of model optimization for poorly studied systems is described in **Note 7** and **Note 8**.

#### 3.5.2 Validation

The final model must be tested on an independent *validation* dataset (*see* **Note 1**). This procedure allows the quantification of the sensitivity of the model, *i*.*e*., to know how well it identifies a secretion system and its different components. However, it is more difficult to assess its specificity, *i*.*e*., its ability to separate true from false instances, since usually there is no reliable information on invalid protein secretion systems.

At the end of the validation process, it might seem tempting (or necessary) to correct the initial model to identify a more significant fraction of the instances of the validation dataset. However, it must be borne in mind that this violates the hypothesis of independence between the model and the validation dataset. Hence, if the model is changed to fit the validation dataset, it can no longer be validated with the same dataset. One can build an additional independent validation dataset to circumvent this difficulty.

#### 3.5.2 Sharing the macsy-models

When satisfied with the generated macsy-model package (systems’ models and corresponding HMM profiles), one might want to share it with the community for easy installation and reuse with MacSyFinder. The macsy-model package installer *macsydata* is distributed with MacSyFinder since v2. It can install macsy-models from a remote repository with the “*macsydata install*” command presented above. But it can also initiate a macsy-model package with the correct file structure using the “*macsydata init*” command and then, upon package completion, check its sanity using the “*macsydata check*” sub-command. Once these steps are passed, the new macsy-model package can be submitted to a dedicated remote repository (the official one is: https://github.com/macsy-models). Once published there, the newly designed macsy-model will be listed and available for automatic remote installation to the community using the “*macsydata available*” and “*macsydata install*” sub-commands. This means that anyone will be able to access and use it. More details can be found in the Modeller guide from MacSyFinder’s documentation.

## 4. Notes

### Note 1. Constructing a reference or validation set of secretion systems

The diversity of the protein secretion systems in the *reference* and *validation* dataset should encompass their diversity in the genomes where they will be searched. Otherwise, the model will miss the instances that are very divergent, in sequence or genetic organization. One can increase the diversity of these sets by sampling instances from all previously described sub-types of a system (*e*.*g*., subtypes SPI1, SPI2, Hrp1, Hrp2, Ysc… for the T3SS [15]). When this information is unavailable, one can use all the known systems to build phylogenetic trees of previously identified key components and then sample representatives of every major clade to build the reference dataset.

### Note 2. Building protein families for the reference secretion systems

We propose the following procedure to build and analyze families of homologous proteins:

1. Store the protein sequences of the reference set of secretion systems in a multi-FASTA file (*e*.*g*., “reference_systems.fasta”). Index the file using the *makeblastdb* command:

~~~
makeblastdb -dbtype prot -in reference_systems.fasta
~~~
2. Run Blastp for each pair of proteins and generate a tabulated output. Pick a statistical threshold of significance for the hits (here E-value < 10^−6^):

~~~
blastp -query reference_systems.fasta -db reference_systems.fasta -evalue 0.000001
-outfmt 6 -out blastall_reference_systems.out
~~~ Alternatively, this can be done using other sequence similarity search tools, such as psiblast or DIAMOND [29, 38].
3. Identify the protein families present in the dataset by clustering the blastp results with Silix [30].

~~~
silix reference_systems.fasta blastall_reference_systems.out -f FAM >
reference_systems.fnodes
~~~
4. Save the different families obtained in separate FASTA files:

~~~
silix-split reference_systems.fasta reference_systems.fnodes
~~~
5. Annotate/label the families. If different components are clustered together, one must disambiguate them. One can split families by clustering the proteins with parameters enforcing higher similarity or alignment coverage. A phylogenetic analysis may also show how to separate two protein families that cluster together (not covered here). Sometimes, one cannot obtain separate protein families (and thus protein profiles). The best solution is to list them all as “exchangeables” under the same single component in the model (*see* **Note 6**).

### Note 3. Design of HMM protein profiles from families of homologous proteins

Follow these steps to build HMM profiles for protein families (*e*.*g*., those obtained following **Note 2**):

1. Store the sequences of each protein family in a different text file in FASTA format. Name the file in accordance with its content, *e*.*g*., “abc.fasta” for the ABC-transporter protein of the T1SS.
2. Make a multiple sequence alignment for each family using MAFFT [25] (or another analogous program):

~~~
mafft abc.fasta > abc.aln-mafft.fasta
~~~
3. Check the quality of each multiple alignments by visualizing them (*e*.*g*., using Seaview [27]). Trim its extremities if they are poorly conserved. Do not remove columns from inside the alignment’s core region, as it will be used to construct the HMM protein profile. Save the edited alignment in a new file, *e*.*g*., “abc.mafft-edit.fasta”.
4. Run *hmmbuild* from the HMMER package [24] on the edited alignment to create the HMM protein profile “abc.hmm”:

~~~
hmmbuild --informat afa --amino abc.hmm abc.mafft-edit.fasta
~~~

### Note 4. Analyzing poorly characterized systems

The typical model-building procedure cannot be used to model systems lacking enough experimentally validated instances to make a reference dataset. In this case, one must use the few known instances to collect homologs from the genome databases by sequence similarity search (*e*.*g*., using blast [23], *see* **Note 2**). Then, the hits can be used to build protein families and HMM profiles. A careful analysis of the components’ co-occurrence patterns usually highlights those that should be specified in the model because they are sufficiently conserved and widespread.

When the reference datasets are small, it is usually better to start building simple models with weak constraints in terms of quorum and co-localization. For example, in a first draft model, every component could be set as a “loner” gene, and the quorum set to two. The model can then be optimized iteratively (*see* section 3.5.1). Naturally, one should be cautious when drawing conclusions from studies using models that could not be validated with an independent dataset of experimentally validated systems.

### Note 5. Setting up protein-specific filtering criteria with the “GA” gathering score in HMM profiles

MacSyFinder v2 can use the GA “Gathering” threshold score in the detection of the proteins with HMM profiles, making protein-specific filtering of the hits possible. The “Gathering” score is defined in HMMER documentation as the minimal score expected for a sequence to be considered part of a sequence cluster or family [24]. The attribution of the GA score can be done by looking at the minimal bit score found for the protein in the detected systems (once the models have been optimized). The information on the minimal bit score can be found in the “best_solution.tsv” output file in the “hit_score” columns. For example, to set up the GA score for the protein PilA of the T4aP after model optimization, the bit scores for all the PilA hits for detected T4aP in a representative genome dataset can be extracted. It can be interesting to analyze the scores distribution shape, to assess the presence of possible outliers and if they could be false positives, etc. A possibility is to evaluate both the distribution of hits within annotated systems and those found outside of annotated systems (*see* **Figure 4** for examples). While the former can be found in the “best_solution.tsv” file (“hit_score” column), the latter can be extracted using the MacSyFinder companion tool *macsyprofile* (see MacSyFinder’s online documentation). Then, the minimum observed value for a PilA component in a detected system can be retrieved from the column “hit_score” and used to establish the GA score in the PilA HMM profile. This can be done programmatically or using a spreadsheet editor. Once the GA score is identified, the HMM profile of the protein (here, “PilA.hmm”) can be opened in a text editor, and the following line added between the “CKSUM” and “STATS LOCAL MSV” lines:

**Figure 4.**
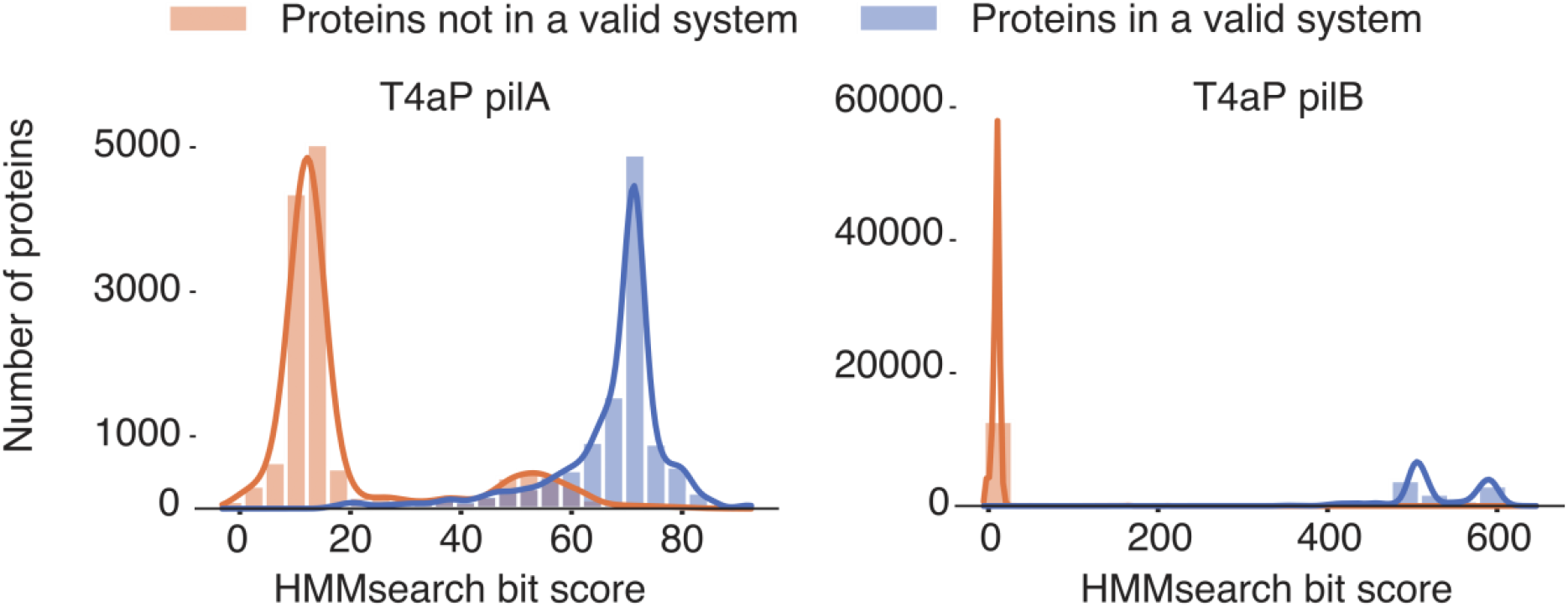
Setting up a GA score threshold for HMM profiles. It is possible to set up a GA “Gathering” score threshold in HMM protein profiles used by MacSyFinder as a way to introduce component-specific filtering of the hits (*see* **Note 5**). This can be done by comparing the bit score distribution for hits found to be part of annotated systems (blue), and for hits not found to be part of systems (orange). This is illustrated in this figure by the case of two proteins from the Type IV a pilus from the TXSScan macsy-model: PilA (left) and PilB (right). We can see that in some cases the two bit score distributions clearly separate from each other (*e*.*g*., for PilB), whereas in some other cases the distributions may overlap (*e*.*g*., for PilA). The GA score threshold can be chosen as the minimal score for a hit found in the system of interest, or any relevant score with respect to the score distribution.

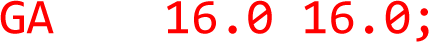

Where 16.0 is the minimal score observed for a PilA protein belonging to an annotated T4aP in the genomes’ annotation results.

The final header of the “PilA.hmm” file will look like this:

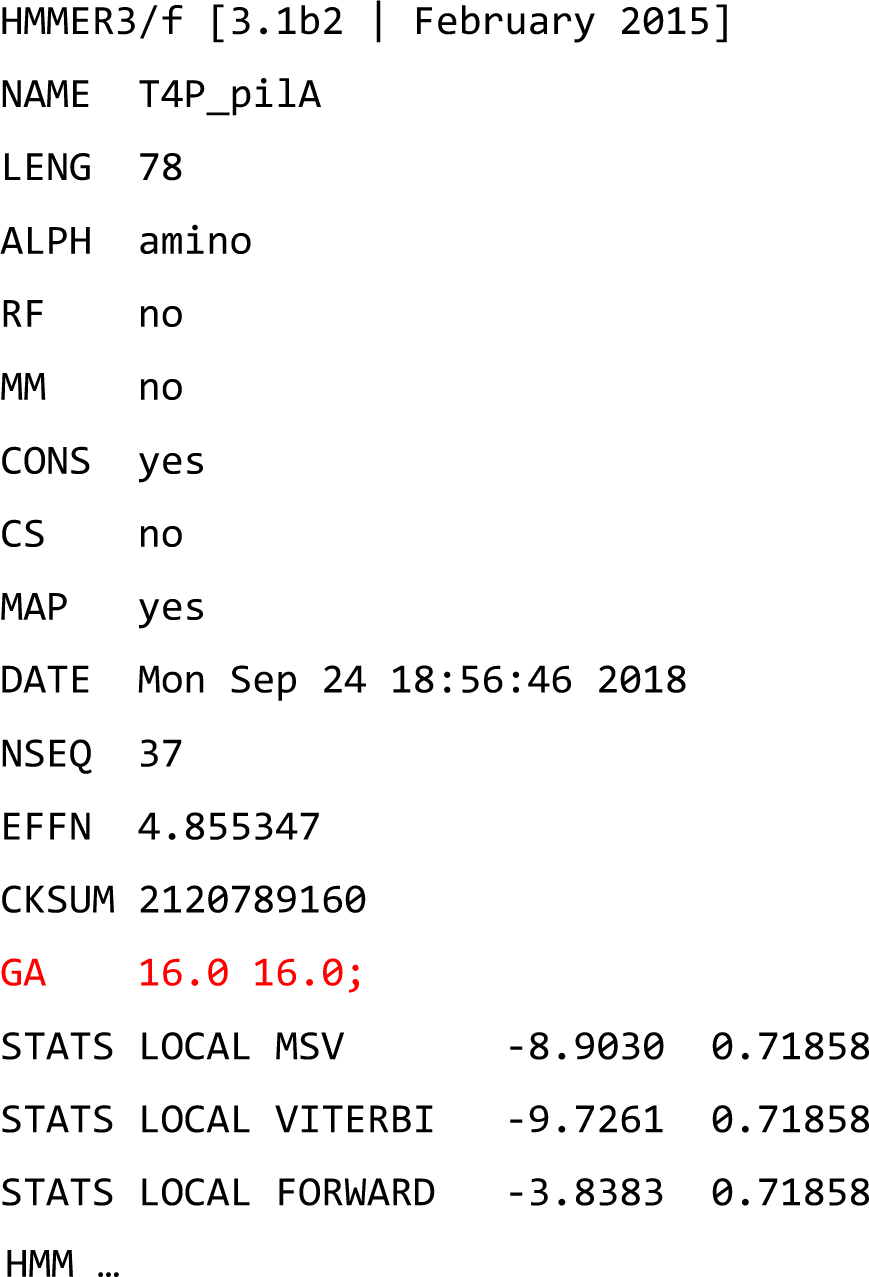

### Note 6. Identification of a component using multiple protein profiles

One single protein profile may not be sufficient to identify all instances of a given component. In such cases, one can associate several protein profiles to a single component in the quorum of the system. These genes should be qualified as “exchangeables”. For example, the T3SS has one of three different sub-families of secretins (T3SS_sctC, T2SS_gspD, Tad_rcpA, depending on the T3SS sub-type [15]). To detect them correctly, one can define them in the T3SS model as follows:

~~~
<gene name=“T3SS_sctC” presence=“mandatory>
 <exchangeables>
<gene name=“T2SS_gspD”/>
<gene name=“Tad_rcpA”/>
 </exchangeables>
</gene>
~~~

With this model, MacSyFinder will identify a T3SS secretin “T3SS_sctC” when it finds a hit for any of these three protein profiles. We show another example with the case of SprA of the T9SS in section 3.2.2.

### Note 7. Optimizing the co-localization criterion

The procedure to optimize the co-localization criterion starts by setting it to a value higher than expected (but not too high, otherwise, several occurrences of the system could be agglomerated). This produces large clusters that are expected to contain all relevant co-localized genes. This parameter may be subsequently refined by plotting the distribution of the maximal distance found between two consecutive components in the clusters of the *reference* dataset. To exemplify, we searched for “single-locus” T6SS in a set of 1528 bacterial genomes using a minimal co-localization distance of 20 genes [10]:

~~~
--db-type gembase --sequence-db bacterial_genomes_proteins.fasta -o
macsyfinder_opt_coloc_T6SS --inter-gene-max-space bacteria/diderm/T6SSi 20 --models
TXSScan bacteria/diderm/T6SSi
~~~

The distribution of distances observed in genomes suggests that a smaller value (*e*.*g*., 14) is enough to identify all relevant clusters (**Figure 5b**).

**Figure 5.**
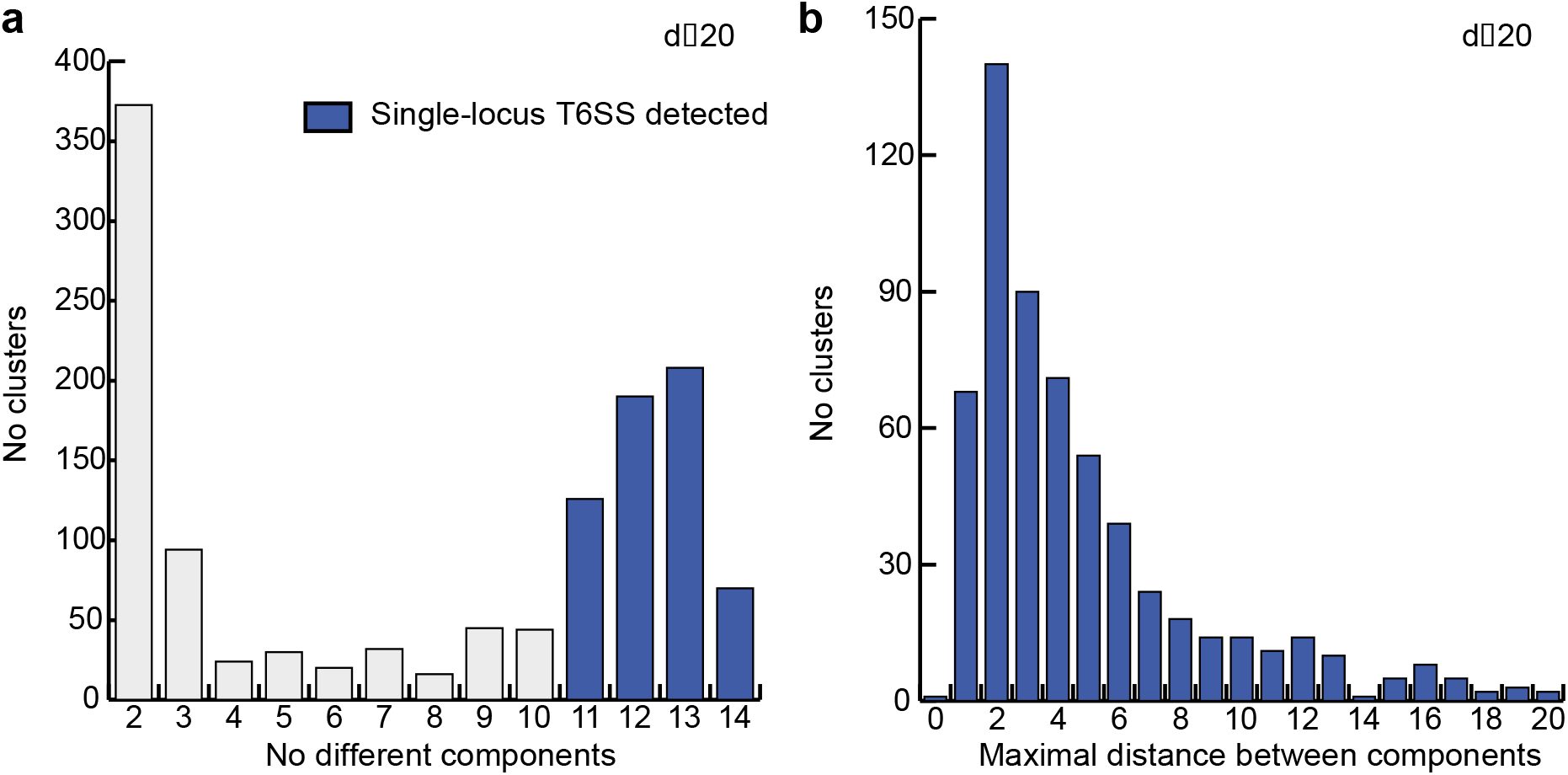
Optimization of the quorum and co-localization parameters for the T6SS^i^. **a**. Distribution of the number of different components of T6SS^i^ co-localized (d≤ 20). **b**. Distribution of the maximal distance between consecutive components in the T6SS^i^ detected with d≤20. The minimal number of components required for a T6SS^i^ was set to 11 in the T6SS^i^ model, which corresponds to the start of the second peak in the distribution represented in panel (**a)**. T6SS^i^ detected as full systems are colored in blue. Figures and legends are freely reproduced with modification from [10] (as specified by the Creative Commons Attribution (CC BY) license version 4.0).

### Note 8. Optimizing the quorum criterion

The quorum can be optimized in multiple ways, depending on the distribution of the secretion systems across genomes.

Suppose the system is encoded at a single locus (but may be found in several copies per replicon). In that case, the quorum can be optimized by studying the distribution of the number of different components detected in each cluster. For this purpose, one can use a model with a very relaxed quorum criterion (*e*.*g*., set to “1”) and draw the distribution of the number of components found in each cluster (with at least one component) (**Figure 5a**). This can be done directly in the command line, for example, using the T6SSi model available in TXSScan:

~~~
macsyfinder --db-type gembase --sequence-db bacterial_genomes_proteins.fasta –o
macsyfinder_opt_quorum_T6SS --min-genes-required bacteria/diderm/T6SSi 1 --min-
mandatory-genes-required bacteria/diderm/T6SSi 1 --models TXSScan
bacteria/diderm/T6SSi
~~~

The number of (accessory and mandatory) components in each cluster can be computed from the easy-to-parse (tabulation-separated) “best_solution.tsv” output file in the ‘macsyfinder_opt_quorum_T6SS’ folder, *e*.*g*., using a Python script based on the *pandas* library (https://pandas.pydata.org) which computes the number of different genes (distinct value in the “hit_gene_ref” column) detected in each different “system/cluster” (distinct value in the “sys_id” column). This specific example shows many clusters with more than 11 components (**Figure 5a**). This is in line with the presence of more than 13 core components in most T6SSi [39]. A final model with a quorum set to 11 would thus accurately identify novel instances of the T6SSi [10].

When the system is typically encoded in a single copy per genome scattered in several loci (“multi_loci”), the analysis of the clusters is less informative, especially if there are other systems in the genome with homologs to these components. Nevertheless, it can be complemented with information on the number of components per replicon. This analysis should use low stringency co-localization and quorum parameters, *e*.*g*., all genes can be set to “loner”, and the quorum set to a small value. We exemplify this by searching to optimize the T9SS model (*see* **Figure 1** and section 3.2.2):

1. Copy the “T9SS.xml” file as “T9SS_loner.xml” in the “./local-TXSScan/TXSScan-refine/definitions” folder, and all the T9SS profiles (“T9SS_*.hmm”) in the “./local-TXSScan/TXSScan-refine/profiles” folder.
2. Add “loner=‘1’” to the definition line of each gene in the new file “T9SS_loner.xml”.
3. Alter the values of the parameters “min_genes_required” and “min_mandatory_genes_required”, either by using the command line (as in the command line below) or by modifying the XML file in the system definition line (“min_mandatory_genes_required=‘1’ min_genes_required=‘1’”).
4. Run MacSyFinder on the modified version of the model:

~~~
macsyfinder --db-type gembase --sequence-db bacterial_genomes_proteins.fasta –o
macsyfinder_opt_quorum_T9SS --min-genes-required T9SS_loner 1 --min-mandatory-
genes-required T9SS_loner 1 --models-dir ./local-TXSScan --models TXSScan-refine
T9SS_loner
~~~

The distribution of “accessory” and “mandatory” components shows many replicons with more than 7 components. The model using a quorum of seven is able to identify novel instances of the T9SS accurately [10].

